# Vorpal: A novel RNA virus feature-extraction algorithm demonstrated through interpretable genotype-to-phenotype linear models

**DOI:** 10.1101/2020.02.28.969782

**Authors:** Phillip Davis, John Bagnoli, David Yarmosh, Alan Shteyman, Lance Presser, Sharon Altmann, Shelton Bradrick, Joseph A. Russell

## Abstract

In the analysis of genomic sequence data, so-called “alignment free” approaches are often selected for their relative speed compared to alignment-based approaches, especially in the application of distance comparisons and taxonomic classification^1,2,3,4^. These methods are typically reliant on excising K-length substrings of the input sequence, called K-mers^5^. In the context of machine learning, K-mer based feature vectors have been used in applications ranging from amplicon sequencing classification to predictive modeling for antimicrobial resistance genes^6,7,8^. This can be seen as an analogy of the “bag-of-words” model successfully employed in natural language processing and computer vision for document and image classification^9,10^. Feature extraction techniques from natural language processing have previously been analogized to genomics data^11^; however, the “bag-of-words” approach is brittle in the RNA virus space due to the high intersequence variance and the exact matching requirement of K-mers. To reconcile the simplicity of “bag-of-words” methods with the complications presented by the intrinsic variance of RNA virus space, a method to resolve the fragility of extracted K-mers in a way that faithfully reflects an underlying biological phenomenon was devised. Our algorithm, *Vorpal*, allows the construction of interpretable linear models with clustered, representative ‘degenerate’ K-mers as the input vector and, through regularization, sparse predictors of binary phenotypes as the output. Here, we demonstrate the utility of *Vorpal* by identifying nucleotide-level genomic motif predictors for binary phenotypes in three separate RNA virus clades; human pathogen vs. non-human pathogen in *Orthocoronavirinae*, hemorrhagic fever causing vs. non-hemorrhagic fever causing in *Ebolavirus*, and human-host vs. non-human host in Influenza A. The capacity of this approach for *in silico* identification of hypotheses which can be validated by direct experimentation, as well as identification of genomic targets for preemptive biosurveillance of emerging viruses, is discussed. The code is available for download at https://github.com/mriglobal/vorpal.

## Feature Extraction Algorithm Overview

In the quasispecies model, the virus organism is represented by the “cloud” of genotypes that can be maintained by the virus within the allowable fitness parameters^12^. In the method proposed here, the frame of reference for the quasispecies “cloud” is reduced to the level of K-length motifs. In order to estimate the connectedness of these K-mers across the input assemblies, a distance matrix between all of the unique K-mers observed across the designated virus genome assemblies is established using hamming distance. Hierarchical clustering is then performed on the resulting distance matrix using an average linkage function, corresponding to the ultrametric assumption used in Unweighted Pair Group Method with Arithmetic Mean (UPGMA) phylogenies, and flat clusters are extracted using a hyperparameter for the distance cutoff of cluster membership. The constituents of these clusters are then aligned and their positional variants represented using the International Union of Pure and Applied Chemistry (IUPAC) nucleic acid notation with degenerate base symbols. These degenerate motifs are mapped back to their respective assemblies. This approach facilitates interpretation of model features in a functional profiling and hypothesis generating context. To demonstrate the effectiveness of this new feature extraction technique, genotype-to-phenotype linear models were trained on various RNA virus groups. A description of the Python implementation of the algorithm is detailed in Methods and the code is available for download at https://github.com/mriglobal/vorpal, along with persistent versions of the models described here-in. A simplified example of the agglomerative clustering step is depicted in Figure 1.

**Figure 1.**
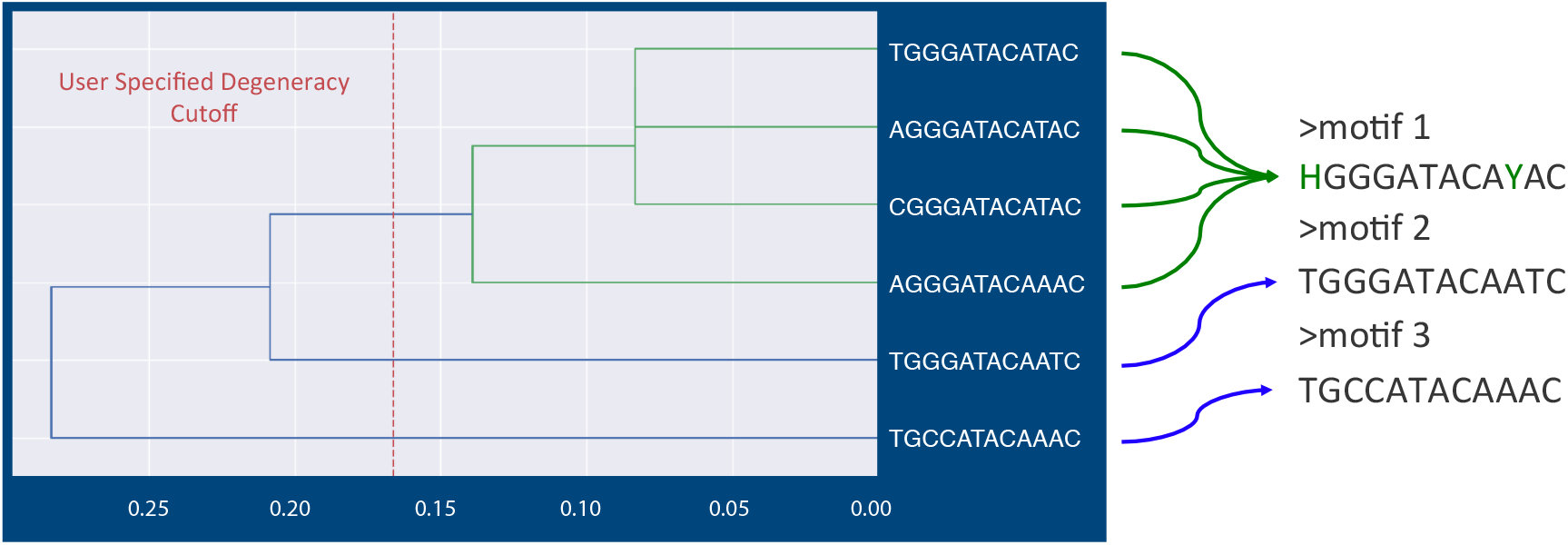
Hierarchical K-mer Clustering. A simplified example of K-mer clustering to produce degenerate motifs. After K-mer counting and filtering on frequency, K-mers are clustered using an average linkage function with hamming distance, or positional agreement, as the metric. The resulting alignments, after tree-cutting at a user specified cutoff, are collapsed into their IUPAC character representation.

By their nature, feature extraction methods make either explicit or implicit hypotheses about what the learner can discover about the data. For instance, in the Natural Language Processing (NLP) domain, the famous “distributional hypothesis” is what forms the theoretical framework for word embedding algorithms such as Word2Vec^13,14^. The hypothesis central to the Vorpal algorithm makes the following predictions about the types of phenomena that could be learned from RNA virus genomics data, if they are relevant to the output label:

1. The predictive motifs are positionally independent
2. The frequency of occurrence of a motif is predictive
3. There are predictive motifs observable only at the nucleic acid level, i.e. in non-coding regions or not observable in the translated product

The strongest predictors for the output phenotypes in the models discussed in this paper demonstrate each of these phenomena.

Three RNA virus groups were chosen to evaluate the methodology, due to their relevance as important human pathogens – Orthocoronavirinae at the sub-family level, Ebolavirus at the genus level, and Influenza A at the species level. The phenotypes for these virus groups were binary output variables corresponding to human pathogen (vs. non-pathogen), human-hemorrhagic-fever-causing (vs. not human-hemorrhagic-fever-causing), and human-host isolate (vs. non-human-host isolate), respectively. The procedure for labeling these phenotypes is detailed in Methods.

This entire algorithm was developed and implemented using Biopython, skbio, and the scipy computing stack contained in the open-source Anaconda Distribution.

## Results

Logistic regression models were fit, in triplicate, for the binary phenotypes described above, across different degeneracy cutoffs for the Ebolavirus and Orthocoronavirinae groups. Due to the training time for the Influenza A models (around 72 hours), instead of exploring different degeneracy cutoffs to find the sparsest feature vector, all Influenza A segment models, which were fit independently, were evaluated with a 1.5 degeneracy cutoff for clustering. Model parameter selection for degeneracy cutoff is visualized in Figure 2. All models were highly accurate on both the training and test sets. Selected models are summarized in Table 1. Construction of the training and test sets is described in the Methods section.

**Figure 2.**
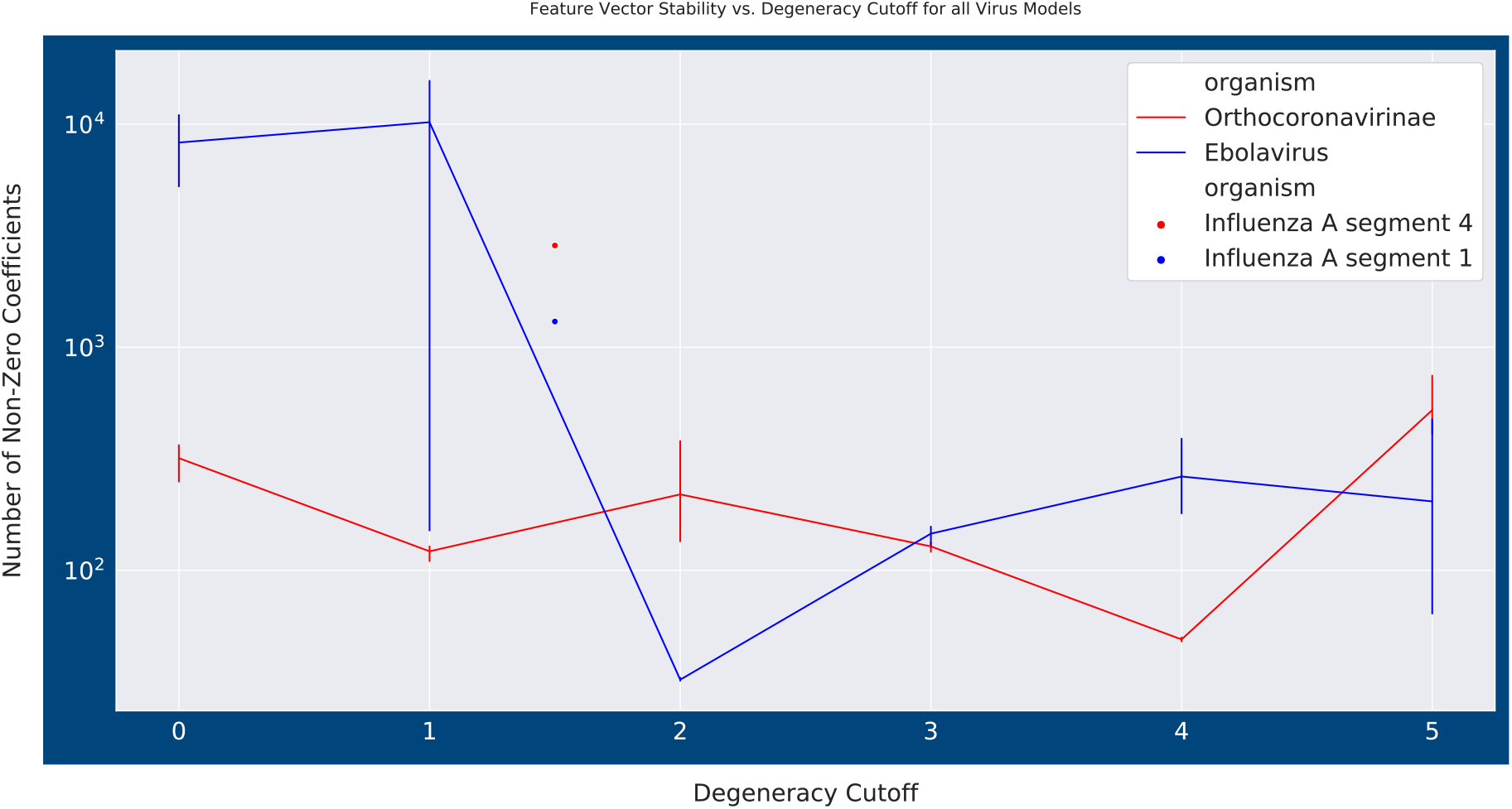
Degeneracy cutoff parameter search. Range of feature vector sizes across different degeneracy cutoff levels. Ebolavirus and Orthocoronavirinae find the least number of non-zero coefficients in the weights vector at 2.0 and 4.0 average degeneracy respectively. They also find very high numerical stability at these cutoffs, with repeated fitting returning almost identical motif set membership. Error bars correspond to standard error of the mean.

**Table 1.** Models Summary. A summary of the attributes for the models built for each RNA virus group that are discussed. NNZs indicate number of non-zero coefficients in the weights vector after regularization.

| Organism | Training Instances (*n*) | Features (*p*) | Model degeneracy cutoff | Quantile | Training Set accuracy | Regularization Method | NNZs | Test Set accuracy |
| --- | --- | --- | --- | --- | --- | --- | --- | --- |
| Orthocoronavirinae | 2278 | 120444 | 4.0 | .95 | 1.0 | LASSO | 48 | 1.0 |
| Ebolavirus | 542 | 92109 | 2.0 | 0.0 | 1.0 | Elastic Net | 33 | 1.0 |
| Influenza A – Segment 1 | 35184 | 11435 | 1.5 | .95 | .9937 | LASSO | 1304 | .9797 |
| Influenza A – Segment 2 | 35252 | 10159 | 1.5 | .95 | .9917 | LASSO | 1330 | .9832 |
| Influenza A – Segment 3 | 35359 | 9985 | 1.5 | .95 | .9969 | LASSO | 1693 | .9769 |
| Influenza A – Segment 4 | 79882 | 41285 | 1.5 | .95 | .9969 | LASSO | 2858 | .9768 |
| Influenza A – Segment 5 | 35492 | 6558 | 1.5 | .95 | .9903 | LASSO | 1104 | .9807 |
| Influenza A – Segment 6 | 57525 | 27435 | 1.5 | .95 | .9897 | LASSO | 1749 | .9833 |
| Influenza A – Segment 7 | 46343 | 3489 | 1.5 | .95 | .9816 | LASSO | 997 | .9759 |
| Influenza A – Segment 8 | 36586 | 5816 | 1.5 | .95 | .9836 | LASSO | 938 | .9802 |

### Explanatory Modeling through Feature Selection

Tables containing the motif identity and corresponding coefficients for the selected models, along with a list of the accession numbers used for training and test sets, are provided as part of the Supplementary materials. We encourage researchers to explore the contents of these models. Below, we analyze a handful of properties of the models to explain their utility in interpretation.

#### Orthocoronavirinae

The model for the Orthocoronavirinae sub-family was built around the phenotype of human pathogen. The motif with the highest coefficient for the human pathogen phenotype, AKRATGKTGTTAATMAA, is an example of the positional independence phenomena that the Vorpal algorithm could learn if it contains information about the response variable. The motif also appears across both Alphacoronavirus and Betacoronavirus group species that infect humans. Interestingly its pattern of appearance in those groups varies in a way not predicated on this taxonomic organization. In the Alphacoronavirus examples that it appears in, namely 229E and NL63, this motif is located in the same reading frame within the spike S2 glycoprotein protein and encodes a conserved QDVVNQ amino acid sequence. However, when it appears in Severe Acute Respiratory Syndrome (SARS), it remains in the same reading frame, coding for a YNVVNK amino acid sequence, but instead occurs in the polyprotein in the N-terminus of non-structural protein (NSP) 15. The other Betacoronavirus member it appears in, OC43, presents this motif in the same reading frame but it has returned to the spike protein as QDGVNK. This motif serves as a signal for human pathogenicity whose importance is based at least partially on its translation, though the domain itself can appear in completely different protein products. It was also recognized that another positive predictor in the model was a motif related to this one, KGATGTTGTTARWCAAY, offset by a single nucleotide. This related motif sometimes co-occurred at the same position as the one mentioned above, and other times appears at a different position in the genome, which suggests this is part of a larger, repetitive motif.

This is summarized in Table 2.

**Table 2.**
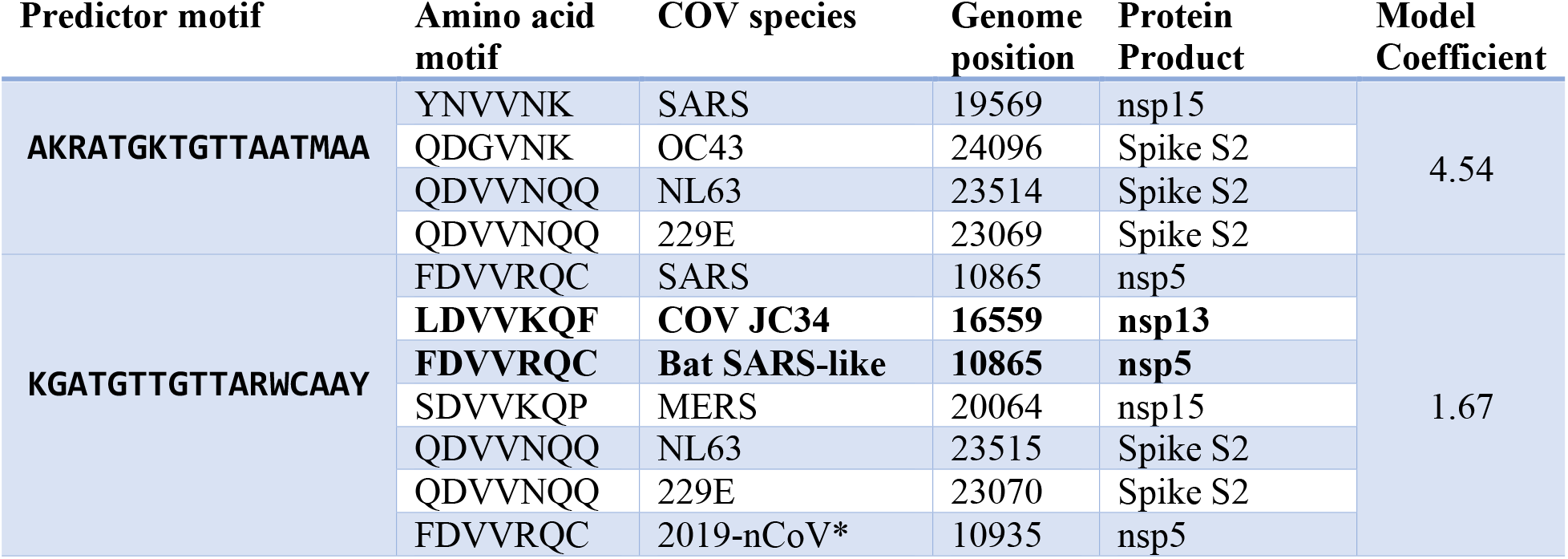
Positive Coefficient Coronavirus (COV) motifs of interest. Organism, genome locations, and corresponding translated products for selected predictors in the Orthocoronavirinae model. Bolded examples are instances labeled Non-human-pathogens in the training set, all others are members of the Human pathogen class. Note: 2019-nCoV was not part of the training set when these models were developed.

#### Ebolavirus

The model for the genus Ebolavirus was specified for a phenotype corresponding to human-hemorrhagic-fever causing, i.e. the African Ebolavirus constituents, and non-human-hemorrhagic-fever causing, i.e. Reston ebolavirus (EBOV). The recently discovered Bombali EBOV, was excluded due to its ambiguity as a human pathogen^15^.

The Ebola model demonstrates the utility of the assumption in the Vorpal algorithm that the feature vector contains information about the frequency of genomic motifs. The preservation of repeated motifs in the 5’ untranslated region (UTR), especially of those in the overlapping UTRs in the Ebola genome, are the predictors of primary importance in differentiating the phenotypes. These repeating motifs, or “motif blocks”, and their corresponding coefficients in the model, are summarized in Table 3 and visualized in Figure 3. These motifs in the 5’ UTRs, specifically in the leading sequence of the L protein, have been previously established as being functionally important to growth kinetics in cell culture^16^. The presence and location of this motif across the Reston and African constituents of the Ebola genus forms an obvious distinguishing factor. The contiguous block of overlapping motifs identified in Table 3, appear across all known Ebolavirus genomes. However, in the Reston version, this block appears only in the 5’ UTR of VP40 and L, which is one of the several genome locations in Reston containing overlapping 3’ and 5’ UTRs. When this motif block occurs in the African-derived constituents of the Ebola genus, it appears in the 5’ UTR of VP40, VP30, and L. The VP40 and VP30 5’ UTRs are characterized by overlapping transcriptional units in African ebolaviruses. In *Zaire ebolavirus,* there is an intergenic region between VP24 and L protein. However, despite the insertion of an intergenic region at this location in *Zaire ebolavirus*, this representation of the motif block is still preserved. Comparison of the transcriptional start and stop signals between Reston and Zaire ebolavirus has been performed before^17^, but the conservation of this motif and this pattern of appearance across the genus has not been established to our knowledge.

**Table 3.**
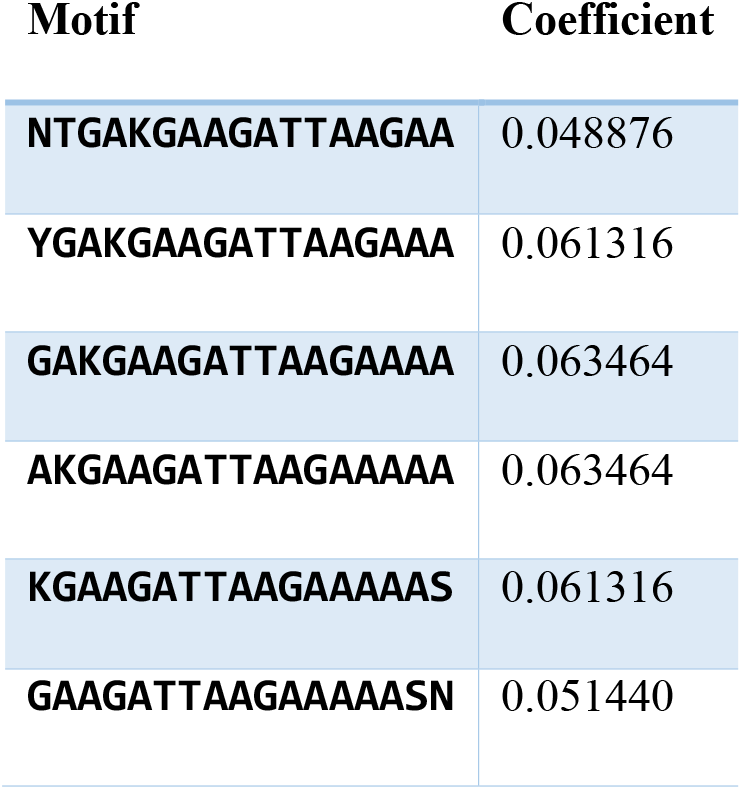
Ebolavirus overlapping UTR “ motif block”. Contiguous motifs that form the 5’UTR overlap conserved at varying frequency across the entire Ebola genus. Identical coefficients represent completely colinear predictors.

**Figure 3.**
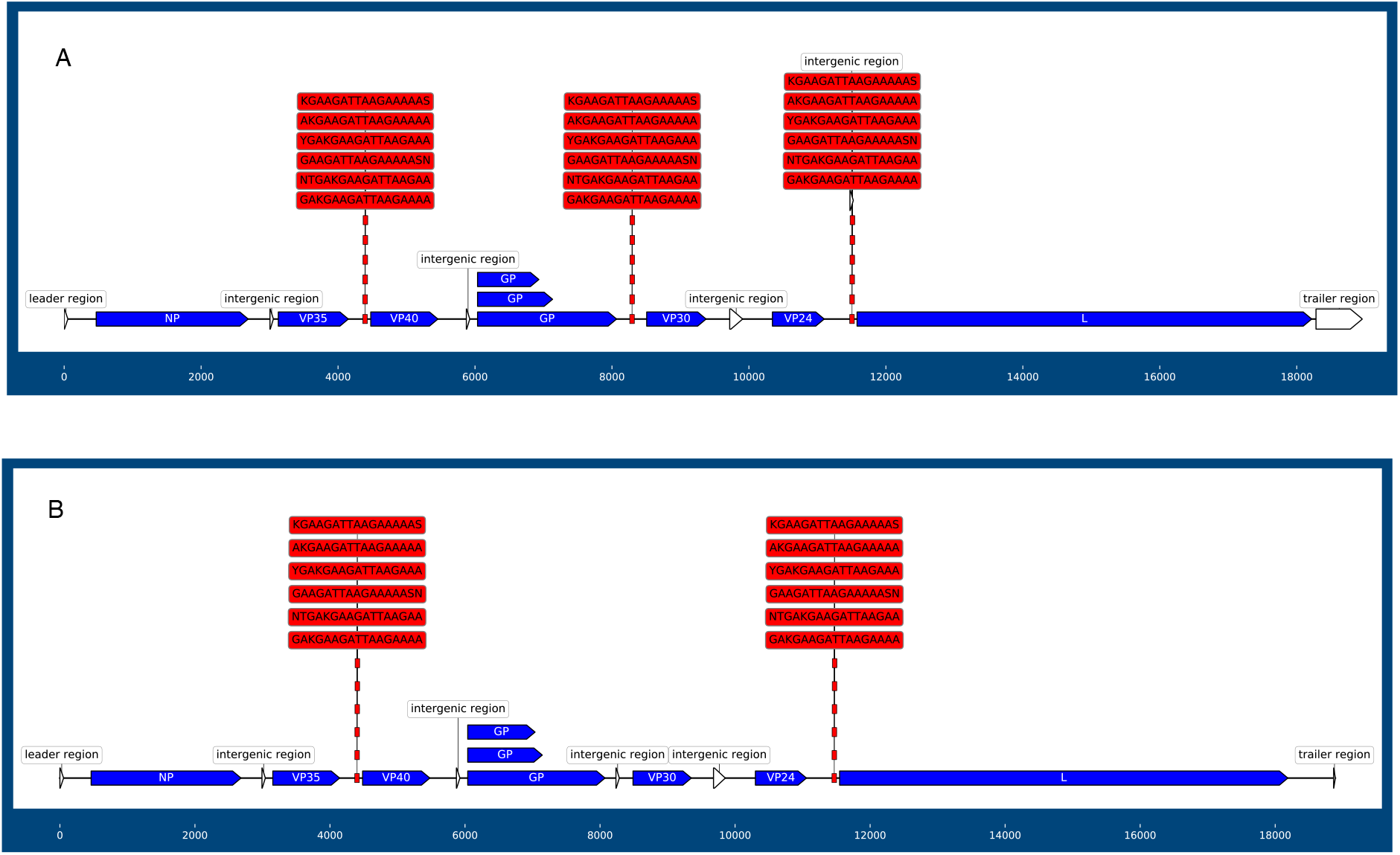
Ebolavirus UTR overlap mapping. Visual comparison of the UTR overlap motifs specified in Table 3. A) Mapping of motifs on the Zaire Ebolavirus genome. The motifs occur three times in the African constituents of Ebolavirus. B) Mapping of motifs on the Reston Ebolavirus genome. The motifs occur only twice in Reston ebolavirus, with the UTR overlap between VP30 and VP24 replaced by an intergenic spacer.

#### Influenza A

The Influenza A model was trained using isolation host as the output variable. As illustrated in Table 1 above, an independent model was built around each segment of Influenza A’s genome. Therefore, the model is trying to find signals of host conformational changes on each segment. However, within the constraints of this paper, only results derived from the segment 4 model will be discussed in detail.

Influenza A’s fourth segment contains the HA gene from which Influenza A strains derive their H subtype designation. In the corresponding model, a pattern was observed in the motif distributions that was common to all of the Influenza A segment models examined. This pattern aligns with the third assumption associated with the Vorpal feature extraction method – some degenerate predictors encode only silent mutations. In other words, the signal for the output label is observed only at the nucleotide level for many explanatory variables. For example, one of the highest coefficient predictors for the human-isolate phenotype, GTCTCTACARTGTAGAA, appeared to be related to one of the motifs amongst the most negative predictors, GGTCTYTACARTGTAGA. These motifs correspond to a location towards the end of the C terminus of the HA2 protein, at the location of a conserved, H1-subtype, N-linked glycosylation site following the transmembrane region^18^. The pattern of appearance for these motifs is described in Table 4A.

**Table 4A.**
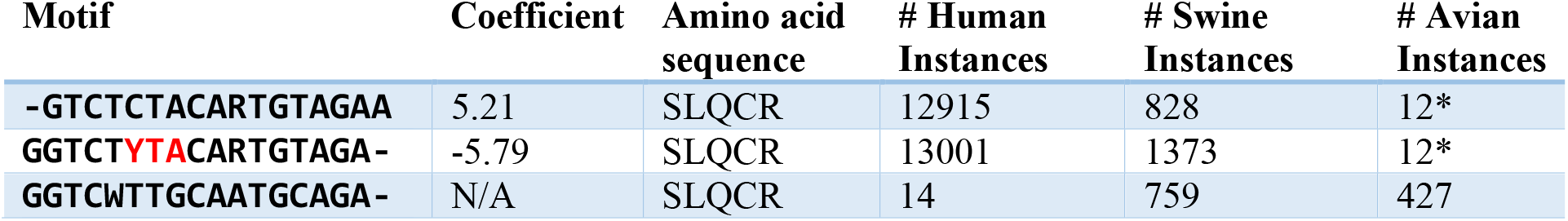
Influenza A HA2 motifs. Shows three overlapping segments where the addition of a degeneracy allowing for the TTA codon for leucine is an important predictor for the non-human conformation for the H1 subtype. The third motif with no coefficient was identified by looking in avian isolates at the same genomic position. This motif was not used by the model but provides additional interpretation of the phenomenon in effect. The only avian flu examples in the model predictors that these motifs appear in are North American Turkey isolates. No other avian examples of any HA gene subtypes contain these motifs utilizing rare leucine codons. This serine at the beginning of this amino acid sequence is the tail constituent of a N-x-S/T glycosylation motif.

Examination of the constituent K-mers of these motifs demonstrated that the allowance of the negative predictor to map to the TTA leucine codon introduced, almost exclusively, swine isolates. The conservation of the CTA leucine codon in the human-isolate predictor is noteworthy because this codon is one of the rare leucine codons in the human genome, with a relative frequency of 7%. Alternatively, the TTA codon being more predictive for swine isolates is notable because while TTA also only has a 7% relative abundance in humans, its abundance in pigs is 6% while the CTA codon is less rare (13% relative abundance)^19^. This mammalian adaptation separates it almost entirely from any avian examples and there appears to be a fitness gradient. When it appears in mammals, there is a higher incidence of the uncommon leucine codon at this location. As previously mentioned, the SLQCR motif is canonical across all H1 subtype examples, including those of chicken and duck. A degenerate motif that mapped to the corresponding position in avian examples was determined to be GGTCWTTGCAATGCAGA. The underlying nucleotide conformations appear to be strictly enforced where the use of the TTG codon for leucine, along with the TGC codon for cysteine, produces 427 avian examples and only 14 human examples. Curiously, the preference for these codons in the avian examples are not correlated with their rarity in those hosts. The TTG codon for leucine has a relative frequency of 13% in mallards, while the CTA and TTA codons are both 6%. A table of the relative codon frequencies by host are noted in Table 4B, and this relationship between motif mapping frequency following a codon rarity gradient in mammals, and the inverse in birds, is visualized in Figure 4.

**Table 4B.** Relative Host Codon Frequencies for HA2 motifs. Shows three overlapping segments where the addition of a degeneracy allowing for the TTA codon for leucine is an important predictor for the non-human conformation for the H1 subtype. The third motif with no coefficient was identified by looking in avian isolates at the same genomic position.

| Organism | Leucine Codons |  |  | Cysteine Codons |  |
| --- | --- | --- | --- | --- | --- |
|  | CTA | TTA | TTG | TGC | TGT |
| Human | 0.07 | 0.07 | 0.13 | 0.55 | 0.45 |
| Avian | 0.06 | 0.06 | 0.12 | 0.6 | 0.4 |
| Swine | 0.13 | 0.06 | 0.1 | 0.61 | 0.39 |

**Figure 4.**
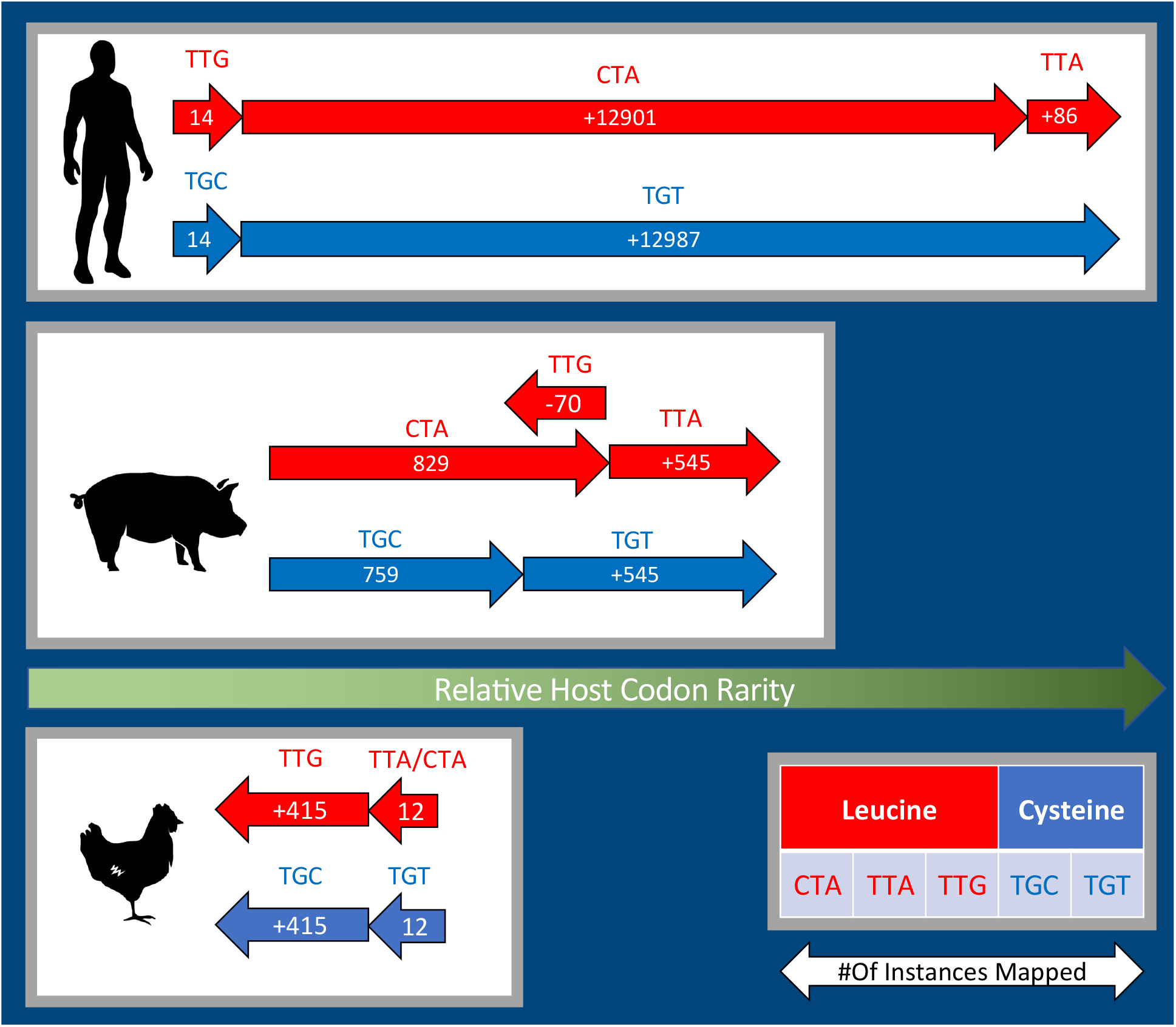
Host Codon Optimizations for H1 subtype. The justification for the coefficients assigned by the model for the motifs specified in Table4A are demonstrated by the clear role that the TTA codon for leucine plays in increasing the probability of a Swine isolate classification for the H1 subtype. Arrows indicating the increase or decrease in total number of motif mappings point in a direction along the relative host codon frequency gradient where rightwards movement indicates optimization towards the lower frequency rank for the corresponding amino acid. Table 4B shows the relative frequencies for these codons across these animal clades. Magnitude of arrows expressing change in number of reference mappings are not drawn to scale.

The predictor variable with the highest coefficient from the segment 4 model is another, more dramatic example, of the phenomenon described above. The identity of the motif, AATGTRACAGTAACACA, and its translated product, NVTVTH, again demonstrate a preference for rare human codons - in this case, valine. Like the example discussed above, this motif is present almost exclusively in human (N=15913) and swine (N=3885) examples of the H1 subtype. The associated NVTVTH amino acid sequence is also completely conserved across all the examples, avian included. The valine codons in the human-isolate versions, almost exclusively GTA, have a relative frequency of only 11% in the human genome. While in the avian examples of segment 4, those codons are switched to GTG, which are the most common Valine codons with a 46% relative frequency in mallards. A motif for the avian version of this was developed using a multiple sequence alignment of the non-human and non-swine isolates of the H1 subtype assemblies in the training set. This motif was established as AAYGTRACYGTGACYCA and mapped back to the training set sequences. When mapped, this new motif was resolved to 480 avian isolates, 33 swine isolates, and nothing else. Unfortunately, unlike the above-mentioned Influenza A motif, the constituent K-mers for this motif were below the quantile cutoff for clustering, and thus, were unable to become a directly observed feature of the model. The use of rare codons, and their tendency to cluster, has been observed across both eukaryotes and prokaryotes^20^. This NVTVTH amino acid motif is also, like the SLQCR sequence described above, an experimentally validated N-linked glycosylation site on the HA gene in H1N1^21^. Rare-codon clusters in association with N-linked glycosylation sites in human pathogens have previously observed in HIV-1 envelope glycoprotein gp120, where the conservation of the rare-codon RNA sequence conferred increased glycosylation efficiency compared to gp120 mutants^22^. Codon optimization efforts for lentivirus envelope protein have also induced non-functional proteins, hypothesized to be related to glycosylation disruption^23^. The fidelity of conformational change in mammalian isolates to these rare codon identities is extremely high. The oscillation between these conformational states is suggestive of another dimension of interpretation that these logistic regression models offer, outside of the examination of the genomic motifs themselves.

### Other Dimensions of Interpretation

The fragility of the phenotype for the Influenza A model resulted in a model with higher complexity than the other RNA viruses studied. However, this provides another avenue for model analysis. Logistic regression classifiers offer not only an output label, but also a probability assignment to the corresponding label. Thus, additional information can be encoded in this output. Figure 5 presents a graphical representation of the distribution of these class probabilities for the training sets for the segments described.

**Figure 5.**
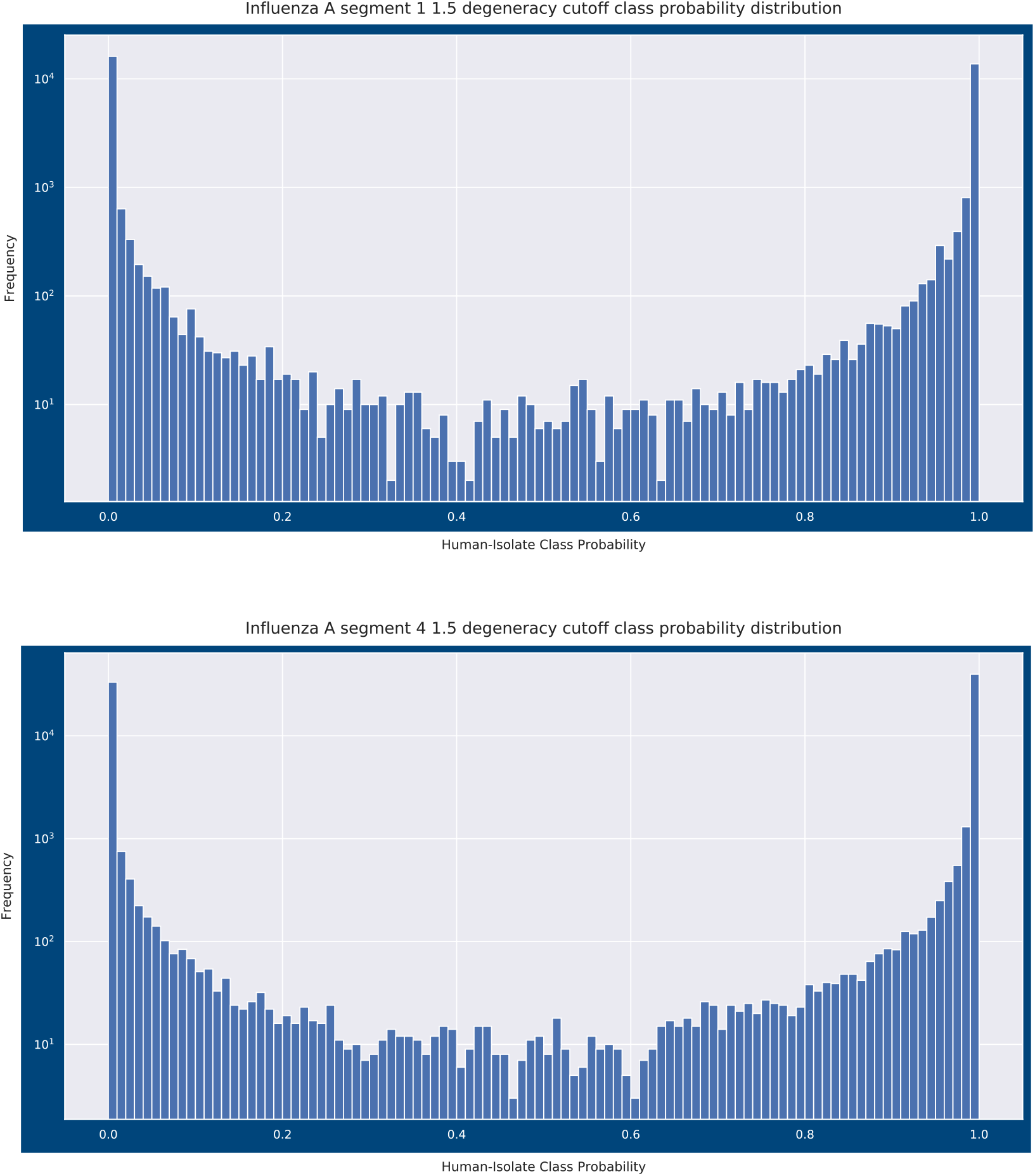
Influenza A training set class probability histograms. The class probability distributions for the Influenza A segment 1 and segment 4 models discussed in this section. The frequency is presented in log scale so it can be observed that the vast majority of class predictions belong to the highest and lowest probability bins. We explore the possibility that instances with class probabilities in the middle of the distribution are in transition between host-isolate states as a results of recent zoonosis events.

The highest coefficient predictor in the model for Influenza A segment 1, which codes for the PB2 polymerase gene, is a motif which represents a mammalian amino acid substitution experimentally observed in a mouse model^24^. This mammalian adaptation was identified as relevant to the temperature sensitivity of the polymerase in H5N1. The reversion of the avian conformation containing the glutamic acid residue, to the mammalian conformation containing lysine, was observed to be approximately six days. By chance, some subset of viral isolates could have been sampled during this window while “in transit” between host-signature genotypes. Thus, the misclassified examples from the training set invite further scrutiny. Of the 79892 instances in the training set, 274 were misclassified, and approximately 10 of these misclassifications were discovered to be mislabeling due to erroneous formatting of the WHO nomenclature. The remainder are examples where the model has, in some cases with a high probability, assigned a classification that disagreed with the class labeling.

One particularly interesting example of this can be seen in a pair of swine isolates (KM289087.1, KM289089.1) misclassified by the model as human, which were attributed to human-to-pig H1N1 transmission events in backyard farms in Peru^25^. A third isolate from this study (KM289088.1) was classified correctly but also expressed some ambiguity in the class designation from the perspective of the class probability. Fortunately, this study included in the publication the sampling dates for the pigs at a central processing facility, allowing the Vorpal algorithm to detect a trend in the data as demonstrated in Table 5. Transition from the human conformation of the virus (from the perspective of the model) to the non-human conformation follows the progression of the calendar date. The original authors had previously speculated about the simultaneous exposure of two of these swine isolates based on phylogeny.

**Table 5.**
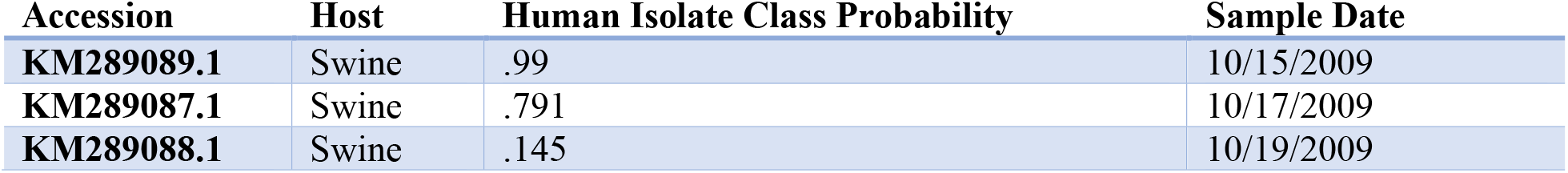
Notable H1N1 Swine Isolates. Transition from the human conformation of the H1N1 virus to the swine conformation from samples in Peru.

| Accession | Host | Human Isolate Class Probability | Sample Date |
| --- | --- | --- | --- |
| KM289089.1 | Swine | .99 | 10/15/2009 |
| KM289087.1 | Swine | .791 | 10/17/2009 |
| KM289088.1 | Swine | .145 | 10/19/2009 |

A second case where a training sample was misclassified as a human isolate from the model was an Influenza A H1N1 instance (KF277197.1) isolated from a giant panda at the Conservation and Research Center for the Giant Panda in Ya’an City, China. There are several plausible hypotheses that could explain the consistent misclassification of this isolate from the model, including the most obvious, that the Giant Panda conformation of the virus is only represented by this distinct example, and thus, the model could not learn the features that may distinguish it from a human-isolated example. However, this assembly was accompanied by a publication which points to a different explanation for model confusion. The paper’s authors, through phylogenetic analysis, suggest that this case was an example of pandemic H1N1 transmitted directly from humans to the pandas^26^. Similar examples are abundant. A pair of misclassified swine isolates were identified in a 2009 publication studying triple-reassortment swine Influenza A infections in people from 2005 – 2009^28^. Both of these human infections were linked to direct contact with sick pigs presented at a county fair within a 3 to 4 day window of sampling. The findings regarding these examples are contained in Tables 6A and 6B. Model prediction probabilities that disagree with the known host source may be useful as a way to infer spill-over events.

**Table 6A.** Midwest, US Influenza A segment four isolates. The local neighbors of A/Ohio/01/2007 (H1N1) and A/Ohio/02/ 2007 (H1N1) identified in Shinde et. al. 2009^27^ as swine influenza virus infections of human hosts at a county fair in 2007. The estimated incubation period for these misclassified training examples was 3-4 days.

| Accession | Year | Location | Human-Isolate Class Probability | Host | Subtype |
| --- | --- | --- | --- | --- | --- |
| **FJ986620.1** | 2007 | Ohio | .420 | Human | H1N1 |
| **FJ986621.1** | 2007 | Ohio | .420 | Human | H1N1 |
| **EU604589.1** | 2007 | Ohio | .310 | Swine | H1N1 |
| **HQ833582.1** | 2007 | Ohio | .069 | Swine | H1N1 |
| **HM461778.1** | 2008 | Ohio | .016 | Swine | H1N1 |
| **HQ378729.1** | 2007 | Iowa | .010 | Swine | H1N1 |

**Table 6B.** Sichuan, CN Influenza A segment four isolates. The Giant Panda isolate misclassified in the training set and its nearest neighbor in the embedding space.

| Accession | Year | Location | Human-Isolate Class Probability | Host | Subtype |
| --- | --- | --- | --- | --- | --- |
| JF277197.1 | 2009 | Sichuan, China | .980 | Giant Panda | H1N1 |
| GQ166223.1 | 2009 | Sichuan, China | .991 | Human | H1N1 |

If the misclassified Giant Panda isolate is observed in context in a two-dimensional embedded space, where the motif feature vectors are used as the input space, then its nearest-neighbor in the lower dimensional representation is a human H1N1 isolate, also from Sichuan, in 2009. In the case of the misclassified swine isolates, they are surrounded in the local neighborhood by H1N1 Swine instances from Ohio and Iowa in 2007 and 2008. This proximity in embedded spaces offers another angle for interpretation, especially in regards to identifying possible spill-over or re-assortment events and is depicted in Figure 6. Neighbors in the local embedding are often temporally and geographically proximal, in addition to sharing host isolate membership. Comingling of class labels in the embedded space potentially offers the opportunity for identification of zoonosis events.

**Figure 6.**
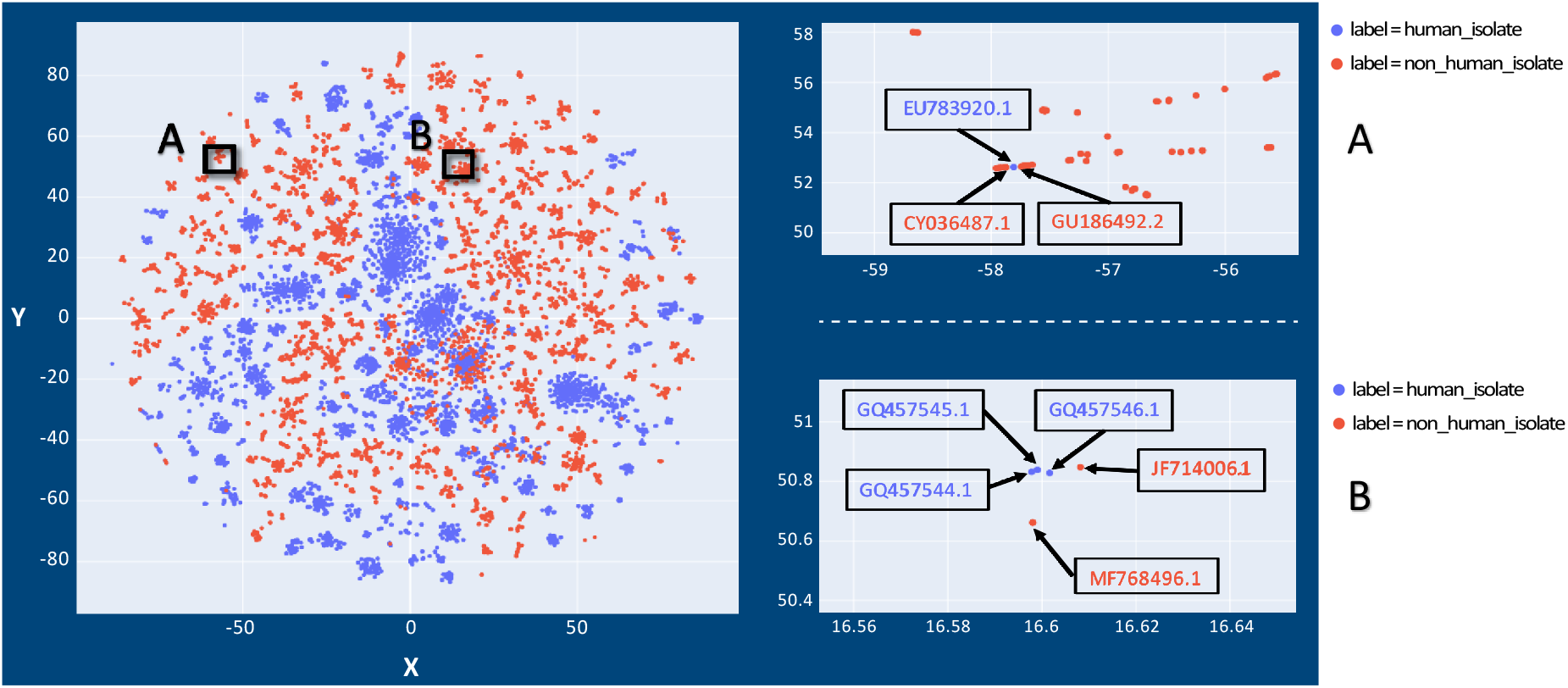
Embedding of Influenza A segment four train set data. Two-dimensional t-SNE embedding of the feature vector for the Influenza A segment four (HA gene). Many clusters can be observed to segregate with respect to the human isolate class label (blue) vs. non-human isolates (red). Close inspection of region (A) identifies linkage of H1N1 isolates from swine and humans likely infected from the same swine population, with the swine-conformation shifting towards a human-conformation. Region A corresponds with data in Table 6A. Close inspection of region (B) identifies linkage of human-conformation H1N1 isolates from humans in Sichuan, China with those from pandas believed to have been infected by direct human contact at a conservation center in the same locale. Region B corresponds with data in Table 6B. Note: Axes in t-SNE plots have no intrinsic meaning except to represent pair-wise distances between points.

Inspection of the embedded space makes it possible to identify candidate events, even if the model has not made a classification error. Examples of these are summarized in Figure 7 and Table 7A and 7B where Influenza A segment 1 (PB2) sequences are embedded into a two-dimensional field. Further experimentation may also help develop models that incorporate a velocity to the conformational changes of host-predictor motifs and estimate temporal distance from a prospective zoonotic event, in a segment-specific manner.

**Table 7A.** New York Influenza A segment one isolates. A human isolate collocated amongst avian isolates in a cluster of H7N2 subtype Influenza examples from New York in 2003. The nearest-neighbor for the human isolate (GU186492.2) is an environmental sample from a live-bird market. The model also encodes the ambiguity of the classification in class probability for the human-isolate phenotype.

| Accession | Location | Subtype | Host | Class Probability | Sample Date |
| --- | --- | --- | --- | --- | --- |
| EU783920.1 | New York | H7N2 | Human | 0.620 | 2003 |
| GU186492.2 | New York | H7N2 | Avian | 0.006 | 2003 |
| CY036487.1 | New York | H7N2 | Avian | 0.013 | 2003 |

**Table 7B.** Saskatchewan, CA Influenza A segment one isolates. A Saskatchewan specific sub-cluster that belongs to a larger cluster of PB2 genes isolated from Swine and co-assorted with H1N1, and H3N2 subtypes in circulation in North America. The human isolates represented in this group belong to pig farm workers who all contracted swine influenza virus, it is presumed, through their place of work^27^. Interestingly, this distinctive genotype of PB2 seems to be preserved across long time frames (2009 to 2015) and is free to re-assort with different Influenza A subtypes (H1N1 and H3N2). In addition, these same isolates had corresponding HA gene sequences published, but the ambiguities seen in the class probabilities for PB2 segment were not observed in the HA gene (i.e. they were all 99% probability human-isolate)

| Accession | Location | Subtype | Host | Class Probability | Onset Date | Sample Date |
| --- | --- | --- | --- | --- | --- | --- |
| GQ457546.1 | Saskatchewan, CA | H1N1 | Human | 0.742 | 6/16/09 | 6/18/09 |
| GQ457544.1 | Saskatchewan, CA | H1N1 | Human | 0.922 | 6/17/09 | 6/19/09 |
| GQ457545.1 | Saskatchewan, CA | H1N1 | Human | 0.769 | 6/15/09 | 6/18/09 |
| JF714006.1 | Saskatchewan, CA | H1N1 | Swine | 0.396 | Unknown | 7/20/09 |
| MF768496.1 | Saskatchewan, CA | H3N2 | Swine | 0.001 | Unknown | 1/15/15 |

**Figure 7.**
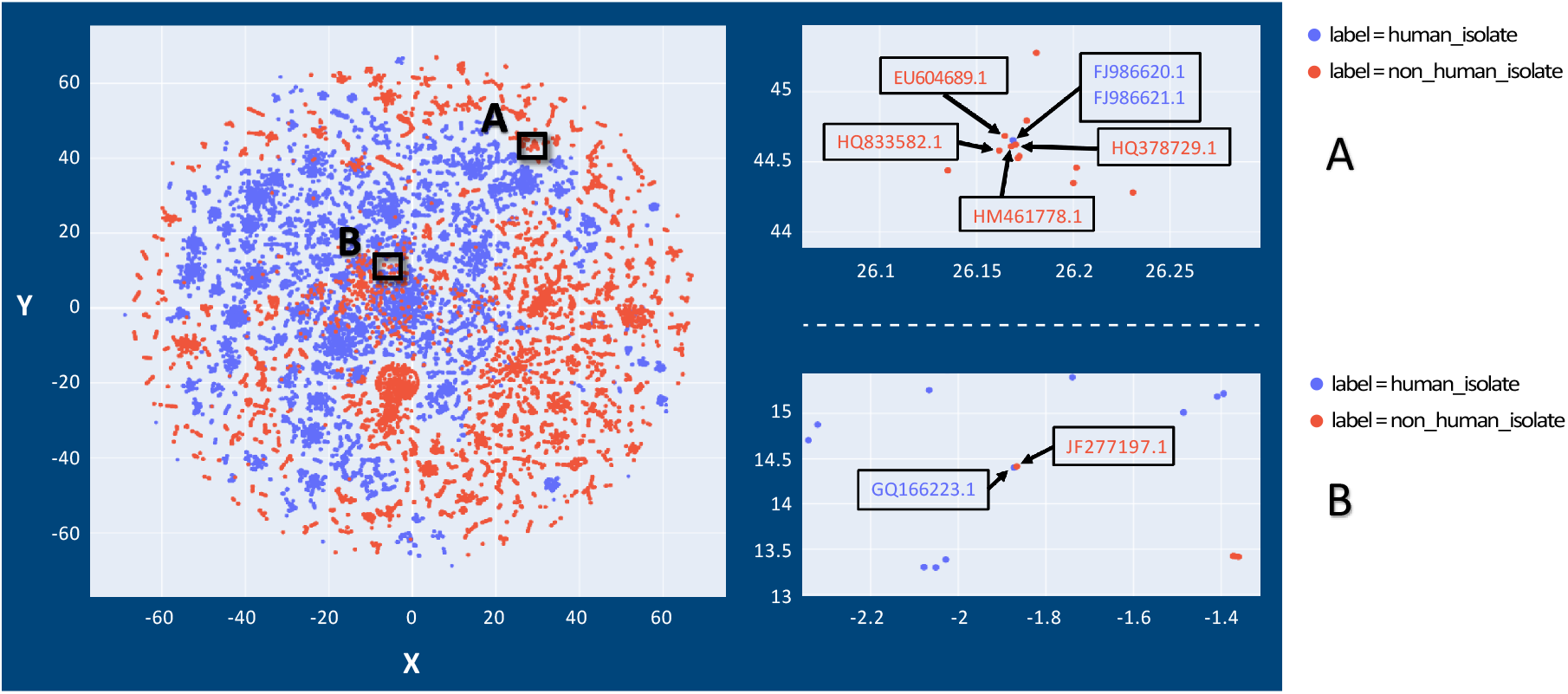
Embedding of Influenza A segment one train set data. Two-dimensional t-SNE embedding of the feature vector for the Influenza A segment one (PB2 gene). Many clusters can be observed to segregate with respect to the human isolate class label (blue) vs. non-human isolates (red). Close inspection of region (A) identifies linkage of H7N2 isolates from birds and a human likely infected from the same avian population at a live bird market within the same locality. Region A corresponds with data in Table 7A. Close inspection of region (B) identifies co-mingling of human-isolated and swine-isolated H1N1 and H3N2 strains from Saskatchewan, Canada. Region B corresponds with data in Table 7B. Note: Axes in t-SNE plots have no intrinsic meaning except to represent pair-wise distances between points.

## Discussion

The observations presented in this paper represent a fraction of the information potentially contained in the developed models using the Vorpal feature extraction algorithm. Efforts to build robust metanalysis tools based on the model outputs is a focus of further development. While we also think the discoveries mentioned herein make a compelling argument for the power of these models in automatically generating hypotheses to direct experiments, we acknowledge the inherent difficulty in leveraging these models for predictive analytics, where, due to the role of evolution, extrapolation to data unsupported at training time is inevitable.

To emphasize the hazard of using these models to predict on new data, the emerging Wuhan pneumonia coronavirus and Bombali ebolavirus provide illustrative examples. The Wuhan COV (MN908947.1) and Bombali ebolavirus (NC_039245.1) assemblies were predicted on using the models denoted in Table 1. The model classified Wuhan COV as 0.004% probable for the Human pathogen phenotype and Bombali ebolavirus as 90.2% probable for the Human-hemorrhagic-fever phenotype. Both of these classifications, especially the Wuhan COV designation, are out-of-step with what is known, or in the case of Bombali, suspected, about these viruses. However, it is possible to imagine these functional profiles leading to a more deterministic understanding of function with which to build a predictive frame work. Nonetheless, improvements in data structure and metadata association may yield better abilities to estimate the probability of future events. Certain observations seen in the models thus far may themselves be predictive of the respective phenotype before it is observed, rather than an effect of it already having occurred. The primary example of this is the predictor identified in the Orthocoronavirinae model. As described in the Methods section, certain assumptions were built into labeling for the human-pathogen phenotype that incorporated theories about the zoonotic provenance of SARS and Middle East Respiratory Syndrome-related (MERS) from civets and camels respectively. Observing human-pathogen predictors occurring in SARS and MERS viruses from non-human hosts could suggest the ability to predict the potential of a virus as a human pathogen in advance of a spill-over event. This is observed in the data. The AKRATGKTGTTAATMAA motif appears in all five of the civet SARS assemblies in the dataset. In the case of the camel isolates, the motif KGATGTTGTTARWCAAY, which is also related to the one mentioned above, is another high coefficient predictor for human pathogenicity and it appears in 231 of the 232 Camel-MERS instances in the training set. This motif also appears in the emerging 2019-nCoV as noted in Table 2.

As for the obstacles for predictive efforts, there are many opportunities for improvements in the collection and annotation of viral genomic data. In Table 1, a slight drift can be observed in the Influenza A model accuracies between the training and test sets. Because the test set represented the most recently isolated viruses, it is attractive to explain this drift as real, i.e. due to evolution. However, there are other factors to control for since the underlying process generating the data has changed over the time period of data collection. The use of cell lines and PCR based amplification of signal for genome assembly, as well as the use of different sequencing technologies suggest other variables to account for. To demonstrated this, a search through the Genbank records for the Influenza A training set members for “passage” annotations revealed that 42.3% of the instances in that set contained such annotations for cell passage. In contrast, the Influenza A test set members, which represents more recently generated data, only contained “passage” annotations in 29.0% of those records.

Lastly, we hope that this analysis demonstrates that the utility of a Global Virome Project is not ambiguous. Controversy about the value of such a project has been described^29^ and this thinking has been reflected in policymakers’ decision to end funding to USAID Predict. If recent estimates of mammalian viral diversity hold true^30^, then marginal increases in monitoring infrastructure combined with new and developing analysis methods, such as Vorpal, might finally deliver the long sought preemptive strategies for emergent diseases, and enable us to more effectively battle those from which we are already suffering.

## Conclusion

The use of this algorithm for genotype-to-phenotype models is just one of the potential applications. Automated molecular assay design and degenerate-motif based phylogenetics are examples of the downstream uses already being investigated. The ability to make use of the latent data that is accumulating in databases, as well as novel surveillance data, is made more tangible with this algorithm. Well-curated and richly annotated metadata promises to allow machine learning and other data science techniques to unleash a torrent of discovery in genomics at large. The mantra we are positing for the infectious and emergent diseases surveillance community is “*More data, Better data, Metadata*.” The techniques to unlock the potential of data-driven genomic science are gathering momentum.

## Methods

### Algorithm

The Vorpal algorithm for feature extraction was developed using the libraries and versions delineated in the requirements.txt document located on the Github. The Vorpal feature extraction algorithm has 3 steps, each corresponding to a script that becomes the Vorpal workflow.

1. kmercountouter_sparse.py

a. Input:

i. a reference genome in FASTA format
ii. a folder containing complete assemblies for the viral group of interest
iii. a parameter for K-mer size
iv. a percent variance argument for filtering out assemblies that are divergent from the reference genome in terms of length
b. Output:

i. a pickled sparse dataframe object containing K-mer counts across every input instance
2. hammingclusters_fast.py

a. Input:

i. A pickled sparse dataframe produced by kmercountouter_sparse.py
ii. The average number of allowed degenerate bases for clustering. This is converted to the equivalent hamming distance cutoff by

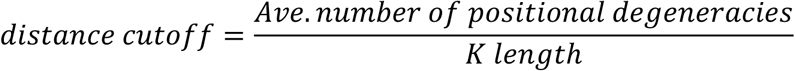
iii. The quantile cutoff for high frequency K-mer filtering
iv. The number of chunks to split the count data into when calculating K-mer frequency. This allows for processing of the K-mer counts table in a memory constrained environment (optional)
v. A temp folder directory to memory map the distance matrix to, again to allow for more memory overhead to be available at the linkage step. (optional)
vi. A memory allocation argument for the development of the distance matrix in chunks. This can be used in conjunction with memory mapping or without it. Uses the sci-kit learn `pairwise_distances_chunked` function instead of the scipy `pdist` function (optional)
b. Output:

i. A multi-FASTA file with degenerate motifs of K length.
3. referencemapping_mp.py

a. Input:

i. A multi-FASTA with all of the assemblies to map to
ii. The multi-FASTA file of degenerate motifs produced by hammingclusters_fast.py
iii. A threads argument for parallel processing
b. Output:

i. A series of BED files with the following column specifications:

| Chr | Start | End | Name | Score |
| --- | --- | --- | --- | --- |
| Accession Number | Start Index | End Index | Motif Identity | $S = \frac{M \text{ instances that motif } i \text{ aligned to}}{\text{Total instances } N} \times 1000$ |

Wrapper scripts for reproducing the models with the parameters described below are also provided as binary_vorpal_model.py and binary_vorpal_model_ElasticNet.py.

### Model Parameters

All models were built around binary output variables using a logistic regression classifier. The models were regularized using either *ℓ*1 or ElasticNet methods, using the liblinear^31^ solver or Stochastic Gradient Descent estimators^32,33,34^ in scikit-learn, respectively. The parameters evaluated for optimization for both approaches were kept uniform for every model fit, with the parameter values searched over listed in the Table 8.

**Table 8.** Grid Search Parameters. Optimization search parameters for regularization methods. Lambda in the LASSO method corresponds to the constraint on the ***ℓ*1** norm of the feature vector while alpha in ElasticNet corresponds to the constraint on the vector magnitude as well as the learning rate for Stochastic Gradient Descent.

| Regularization | Term | Search values | Cross<br>Validation Folds |
| --- | --- | --- | --- |
| **LASSO** | lambda | 1.0e<sup>-4</sup>, 7.742e<sup>-4</sup>, 5.995e<sup>-3</sup>, 4.642e<sup>-2</sup>, 3.594e<sup>-1</sup>, 2.783, 2.154e<sup>1</sup>,<br>1.668e<sup>2</sup>, 1.292e<sup>3</sup>, 1.0e<sup>4</sup> | 5 |
| **ElasticNet (.15**<br>**ℓ1 ratio)** | alpha | 1.0e<sup>-1</sup>, 1.0e<sup>-2</sup>, 1.0e<sup>-3</sup>, 1.0e<sup>-4</sup>, 1.0e<sup>-5</sup>, 1.0e<sup>-6</sup> | 5 |

All of the input parameters for feature extraction and the rationale behind the use and tuning of each parameter, and their relation to the corresponding model discussed above is provided here.

### Feature Extraction Parameters

The first parameter, K length, can be a variable input, but in the development of these methods was fixed at 17. The decision to set the k-length at 17 had many facets. The first is that the feature space should be large enough, that the introduction of degenerate positions does not cause a complete collapse of feature structures. Evaluation of optimal K length for specific tasks has been performed in many contexts. For phylogenetic representations of viruses, an optimal range of 9 to 13 has been proposed^35^, for the optimal uniqueness ratio in plant genomes a K length of 20 has been identified^36^, and in phenetic analysis of bacteria a K length of 31 has been demonstrated to yield the best balance between sensitivity and specificity in intra- and interspecies distance analysis^4^. However, defining a subspace that lends itself to genotype-to-phenotype model interpretability should have the following desiderata:

1. K-size motifs should map to mostly unique genomic loci. In other words, sparsity in the weights vector is influenced by sparsity in the input vector.
2. K-size should be small enough, that the feature space inflation is not catastrophic to memory constraints.

**Figure 8.**
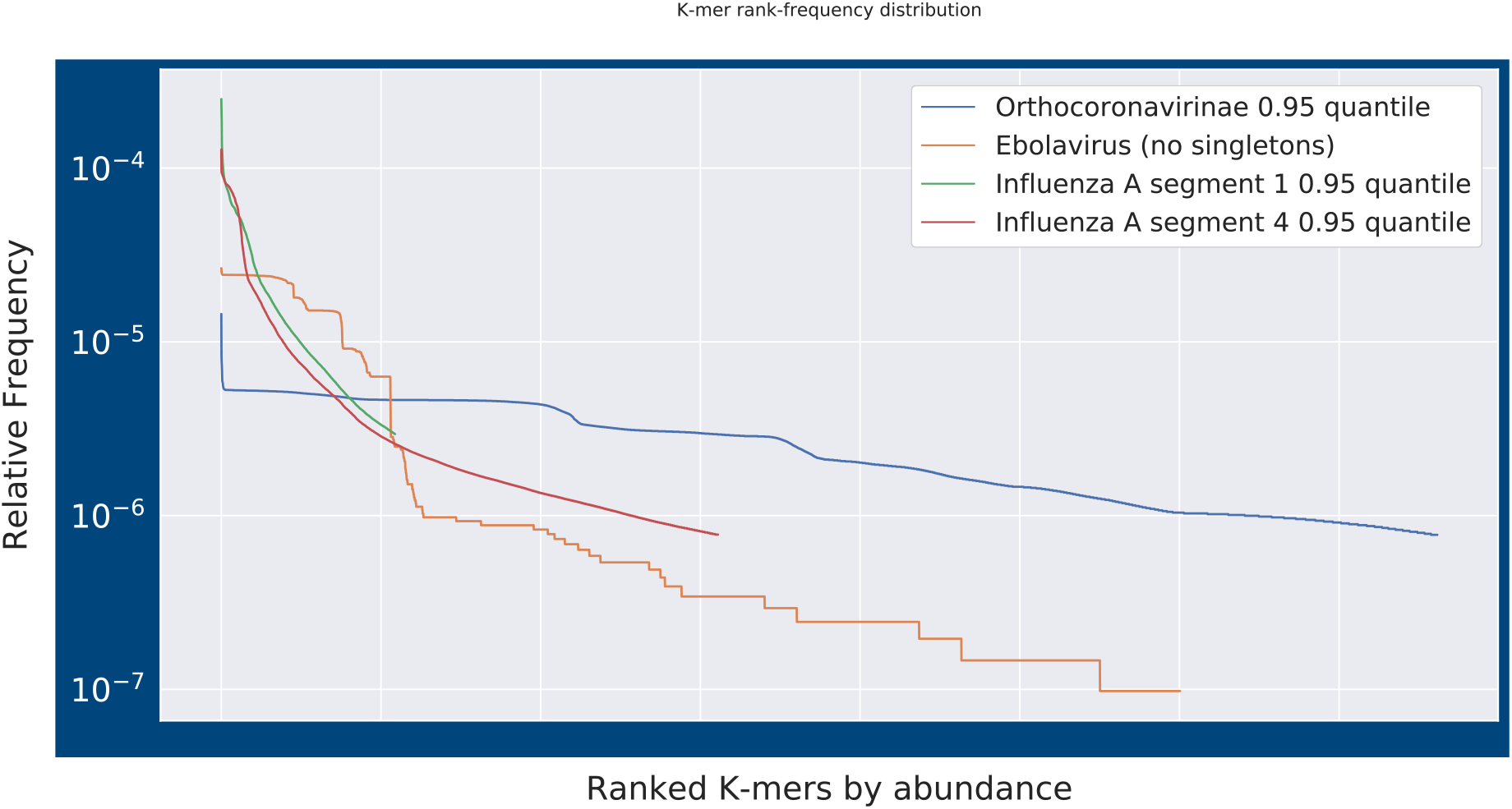
Post-quantile filtering K-mer distribution. K-mer rank-frequency distribution plots for Influenza segments 1 and 4, Ebolavirus, and Orthocoronavirinae at the quantile used in the models discussed. Frequency is calculated as number of instances the ranked K-mer appears in.

This method implements canonical K-mer counting, where the reverse complement of a K-mer is counted as the same time as the forward oriented K-mer, because of uncertainty about strand orientation in the input data. It was known that there were example assemblies in GenBank for Lassa virus where different instances had inconsistent strand reporting. This assumption seems to be unwarranted for the viruses selected for this study and could be removed for future implementations. It should be pointed out that, while maintaining this assumption seems wasteful from a memory overhead perspective, certain features could only be revealed through this canonical approach, such as hair-pin complements in RNA secondary structures, where the resolution of this structural motif is only possible when compared to the K-mer produced by the complementary region. Other dimensionality reduction techniques, namely high-frequency K-mer filtering, allowing the feature extraction to remain tractable, given the computing resources available for this study. The effect of this canonical approach, and the information it potentially encodes in the feature space, is demonstrated in Table 9A and 9B.

**Table 9A.** Orthocoronavirinae Feature Extraction Summary (0.95 quantile). Summary statistics for feature extraction for Orthocoronavirinae from 0.0 to 5.0 degeneracy cutoff for clustering. Feature space tracks the dimensionality reduction introduced by degeneracy to motifs that map back to training set. Initially, since no odd-length K-mer can be a reverse complement of itself, canonical K-mers counted compared to those mapped should be half. As degeneracy is introduced, the Motif/Feature ratio is expected to converge to 1.0, which describes a single motif of all “N” symbols. This ratio tracks the amount of previously distinct motifs now represented as a single feature. The final column, shows the phenomena of motifs that are now reverse complements of themselves as a result of degeneracy, contributing to the inflation of the Motif/Feature ratio. Of note in the Orthocoronavirinae features, is while dimensionality reduction continues with the allowance of more degeneracy, the fraction of those resulting features that have corresponding reverse complements in the feature set does not increase past 4.0 degeneracy.

| Degeneracy Cutoff | # Motifs | Average degenerate positions | Average degenerate bases represented | Feature space | Motif/Feature ratio | Reverse Complement Motif Ratio |
| --- | --- | --- | --- | --- | --- | --- |
| 0.0 | 380726 | 0 | 0 | 190363 | 0.5 | 0.00177 |
| 1.0 | 354138 | 0.074744873 | 0.14982295 | 177375 | 0.50086407 | 0.00339 |
| 2.0 | 331512 | 0.20877374 | 0.419245759 | 167825 | 0.506241101 | 0.02309 |
| 3.0 | 266349 | 0.939684399 | 1.894649501 | 146501 | 0.550033978 | 0.13751 |
| 4.0 | 183004 | 2.855790037 | 5.835227645 | 120444 | 0.658149549 | 0.26588 |
| 5.0 | 150418 | 3.787837892 | 7.956893457 | 106462 | 0.707774336 | 0.24210 |

**Table 9B.** Ebolavirus Feature Extraction Summary (0.0 quantile). Summary statistics for feature extraction for Ebolavirus from 0.0 to 5.0 degeneracy cutoff for clustering. A larger fraction of the features at the highest degeneracy allowance produced contain corresponding reverse complement motifs in the feature set in the Ebolavirus data than in the Orthocoronavirinae data. This could be attributable to the high frequency K-mer quantile cutoff utilized in the Coronavirus group, or it could allude to generally higher fraction of the Ebolavirus genomes having self-complementation than Coronavirus genomes.

| Degeneracy Cutoff | # Motifs | Average degenerate positions | Average degenerate bases represented | Feature space | Motif/Feature ratio | Reverse Complement Motif Ratio |
| --- | --- | --- | --- | --- | --- | --- |
| 0.0 | 300196 | 0 | 0 | 150098 | 0.5 | 0.00052 |
| 1.0 | 227131 | 0.316856792 | 0.638543396 | 113622 | 0.500248755 | 0.00114 |
| 2.0 | 183592 | 0.692203364 | 1.401003312 | 92109 | 0.501704867 | 0.00620 |
| 3.0 | 151326 | 1.265717722 | 2.599824221 | 77855 | 0.514485283 | 0.04837 |
| 4.0 | 108073 | 2.715136991 | 5.716635978 | 61648 | 0.570429247 | 0.18857 |
| 5.0 | 86031 | 3.884692727 | 8.358347572 | 53537 | 0.622298939 | 0.27256 |

**Figure 9A.**
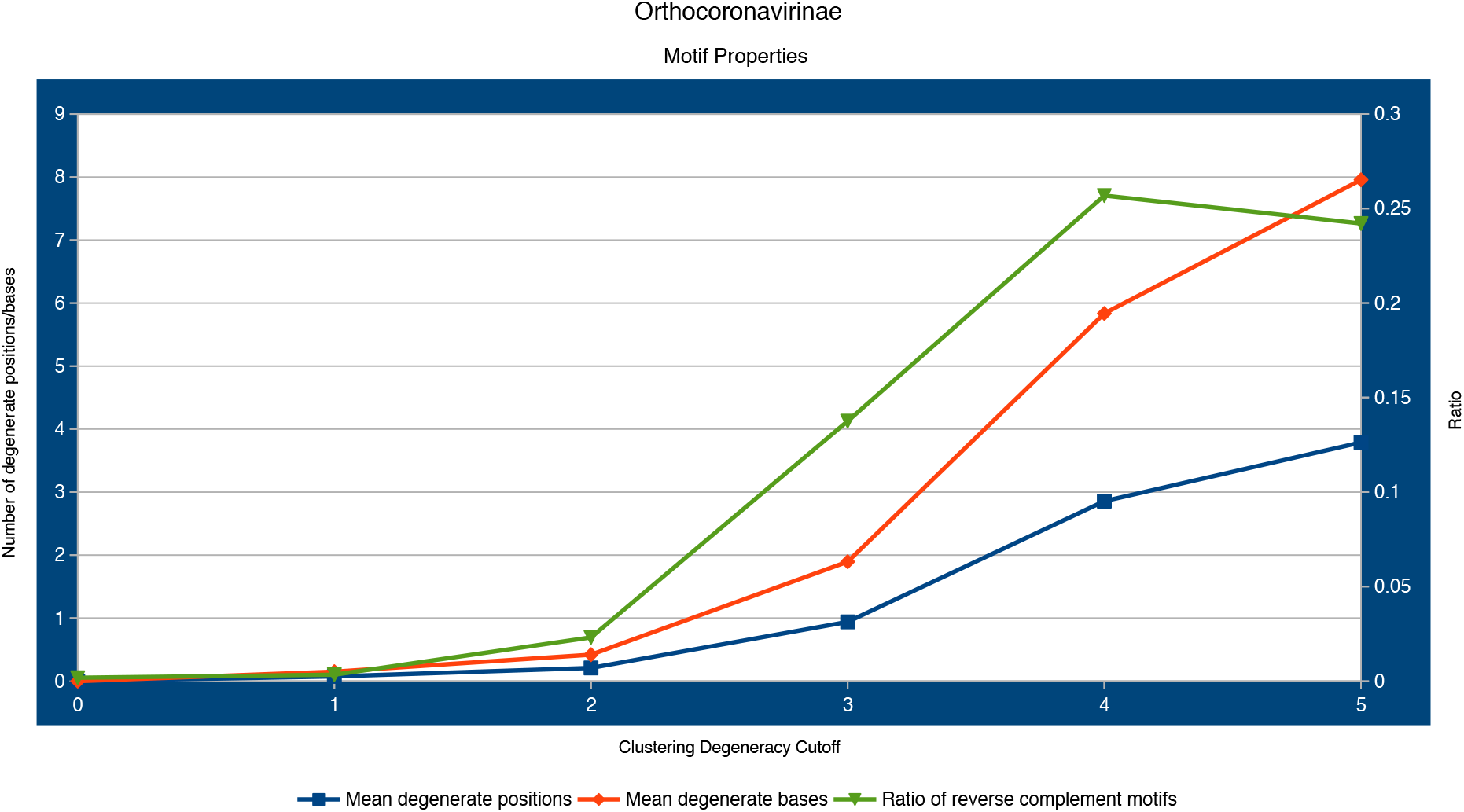
Line plot for three of the selected columns from Table 10A. The plateau reached at 4.0 degeneracy for the ratio of reverse complement motifs is clearly evident.

**Figure 9B.**
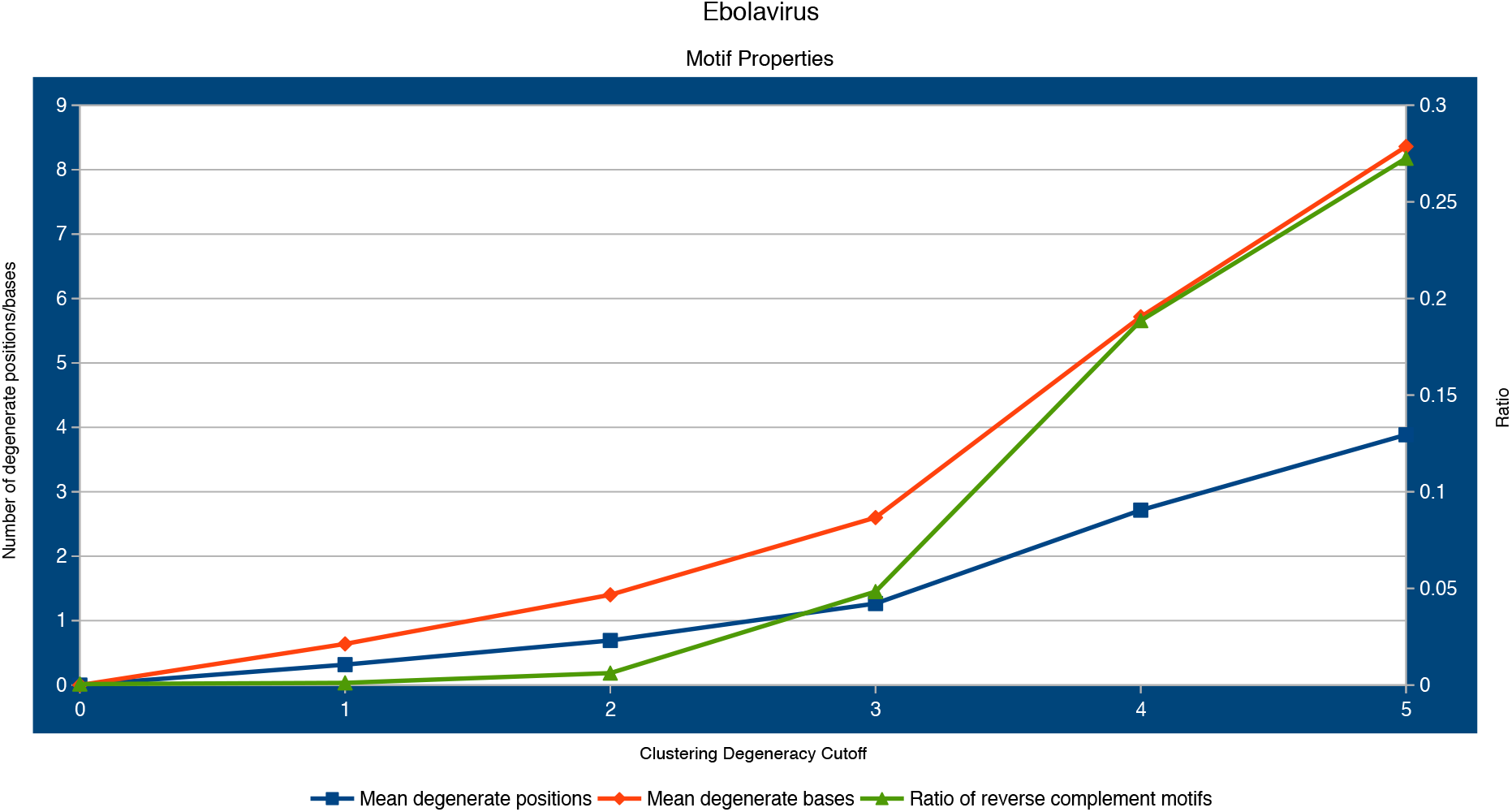
Line plot for three of the selected columns from Table 9b. The relationship between mean number of degenerate positions in the motifs and the mean number of degenerate bases represented is very similar between Ebolavirus and Coronavirus.

Applying a filter to the K-mers that are allowed to proceed to the clustering step has two purposes. The first is to denoise the data by removing low abundance features that could be the result of error or other transient sources of variance. The removal of these K-mers is achieved through a parameter specified at the clustering stage, the K-mer quantile. Singletons, or K-mer that are unique to a single instance, are always removed no matter the quantile specified. It was discovered that allowing the singletons to form motifs through agglomerative clustering introduced instability into the model parameter estimation (data not shown). Contribution to frequency is determined not by cumulative sum of count across every instance but rather frequency of presence across the sample instances. This is identical to the way the “TopN” score is calculated for K-mers in PriMux primer design software^37^. Using a K-mer frequency filter selects for a conserved variance signal. This is a reasonable heuristic to introduce, especially for predictive models, where these high-frequency K-mer derived motifs are the those with the presumed highest probability of appearing in a novel example of a related organism in nature. The second function of this feature extraction parameter, made reference to above, is as a dimensionality reduction technique to make K-mer clustering more tractable in the current algorithm implementation, given limitations in computational resources. Memory constraints during the tree building step represents the primary bottleneck with the scipy implementation of the nearest-neighbors chain algorithm for average linkage using 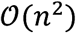 memory^38,39^. The user specifies an average number of degenerate bases to apply when flat clustering. This number is then divided by the K length specified to estimate the corresponding hamming distance to provide as the max distance for flat clustering. After flat clusters are grouped into alignments and a degenerate motif of the alignment is generated by collapsing each position in the K length alignment into the IUPAC symbol matching the bases seen at that position. This clustering of K-mers, and subsequent representation as degenerate motifs, is another layer of dimensionality reduction similar to lemmatization of words in a Natural Language Processing (NLP) feature extraction technique^40^. Much of this approach could be described as modifications of equivalent NLP feature extraction and modeling strategies. It should be noted however, that data preparation techniques such as term frequency-inverse document frequency (tf-idv), were considered inappropriate to apply in this circumstance for multiple reasons. First, “document” length was invariant in the sense that complete assemblies were the only instances allowed in the training data, and differences in genome sizes within the taxonomies considered were considered irrelevant. Second, document terms, in this case K-mer motifs, that follow a frequency pattern similar to the word “the” in the English language are not present. Additionally, for this reason, the data was not normalized, however to improve convergence speed this could be a future improvement.

### Phenotype Labeling

Phenotype labels for the different organisms modeled were applied using a variety of strategies with some specific assumptions introduced for labeling of the Orthocoronavirinae group. In the cases of Ebolavirus and Coronavirus, taxonomy was used as a guide for phenotype labels, where knowledge about the phenotype of interest was usually easily delineated along taxonomic boundaries. For Influenza A, the World Health Organization nomenclature for Influenza strain identification, which is encoded in the FASTA header, was parsed for labeling of human isolate^41^. For those FASTA headers which contained malformed strain identifiers, an ambiguous labeling was applied and removed from the training set.

The following explicit assumptions were applied when labeling viral instances for the human pathogen phenotype in the Orthocoronavirinae model. First, since most transmissions of Middle East Respiratory Syndrome-related (MERS) betacoronavirus to humans have been zoonic events traced to dromedary camels, all camel isolates for MERS coronavirus were labeled in the positive class corresponding to human pathogen. Likewise, in the cases for Severe Acute Respiratory Syndrome-related betacoronavirus, since the initial outbreak had been theorized to begin from a zoonic event from infected palm civets at a market in Guangdong, China, along with a specific civet spill over event documented in a waitress and a customer in a restaurant in Guangzhou^42^, all civet SARS-like isolates were also labeled as belonging to the positive class. However, since there is no clear evidence of bat-Coronavirus-to-human transmissions, the assumption was built-in that bat isolates of both MERS-like and SARS-like betacoronaviruses were not part of the human pathogen class. In the instance of MERS-like bat isolates, examples have been found across wide geographic ranges, such as South Africa, while human cases appear to restricted to areas where Saudi Arabian dromedary camels are present^43^ or hospital acquired infections. The same is true of SARS-like bat isolates discovered in caves in China, where assemblies from these isolates show varying similarities to the strain from the 2003-2004 outbreak but not the sum of them^44^.

Training sets were developed from the un-clustered Reference Viral Database^45^ (RVDB) version 14 published October 1^st^, 2018. Accessions for designated taxonomic groups were derived from National Center for Biotechnology Virus^46^ and then used to extract the associated assemblies from RVDB. Test sets were developed from RVDB version 15 published February 6^th^, 2019 using the references for the modeled organisms that had been added between version releases.

### Embedding Visualization

The same feature vectors used to produce the logistic regression models were topic modeled similarly to Latent Semantic Analysis^47^ (LSA) using a truncated Singular Value Decomposition (SVD) to a 500 component subspace, which was then subjected to a t-distributed Stochastic Neighbor Embedding^48,49^ (t-SNE) to a two-dimensional space to observe the local structure of the Influenza A viral assemblies. Both of these methods were employed using the associated classes in Scikit-learn. Visualization and exploration of the embedded space was facilitated by Plotly^50^.

